# Visualization and estimation of stroke infarct volumes in rodents

**DOI:** 10.1101/2023.07.14.547245

**Authors:** Rebecca Z. Weber, Davide Bernardoni, Nora H. Rentsch, Beatriz Achón Buil, Stefanie Halliday, Mark-Aurel Augath, Daniel Razansky, Christian Tackenberg, Ruslan Rust

**Author notes:** **Correspondence** Ruslan Rust.

## Abstract

Stroke volume is a key determinant of infarct severity and an important metric treatments evaluation. However, accurate estimation of stroke volume can be challenging, due to the often confined 2-dimensional nature of available data. Here, we introduce a comprehensive semi-automated toolkit to reliably estimate stroke volumes based on (1) whole brains *ex-vivo* magnetic resonance imaging (MRI) and (2) brain sections that underwent immunofluorescence staining. We located and quantified infarct areas from MRI three days (acute) and 28 days (chronic) after photothrombotic stroke induction in whole mouse brains. MRI Results were compared with measures obtained from immunofluorescent histologic sections of the same brains. Using our toolkit, we found that infarct volume determined by post-mortem MRI was highly correlated with a deviation of only 6.6% (acute) and 4.9% (chronic) to the measurements as determined in the histological brain sections indicating that both methods are capable of accurately assessing brain tissue damage.

## Introduction

Stroke continues to be a major cause of mortality worldwide and remains the primary cause of acquired disability, resulting in significant personal, social, and economic burdens. Although many drugs and therapies have been tested over the last decades, there is no effective strategy yet to completely prevent or cure the disease.^1^ Thrombolytic therapy using recombinant tissue-type plasminogen activator (rtPA) is the only approved drug for clinical use during the acute phase of ischemic stroke.^2^ Its application, however, must be administered within 4.5 hours of symptom onset to be effective, limiting the options available to patients who are unable to seek medical attention quickly. Ergo, there is an urgent need for developing new medical treatments for acute ischemic stroke. Yet, translation of preclinical stroke research is rare, which may be attributable to the lack of standardization in experimental protocols, making it difficult to compare results across studies.^3^

Rodent models have long been employed as one of the important methods for understanding ischemic stroke mechanisms. In many preclinical stroke studies, treatment efficacy is assessed through statistical comparison of infarct sizes of treatment versus placebo groups. Several methods are suitable to investigate the extent of brain damage in experimental stroke models. New nanotechnology-based techniques such as magnetic particle imaging (MPI) or the use of tracers (PET, SPECT) are becoming more common in preclinical research.^4^ Magnetic resonance imaging (MRI) – a technique routinely applied in humans – can also be used to define ischemic lesions in rodents *in* and *ex vivo*. Nevertheless, traditional approaches using post-mortem histological examination to study the extent of neuronal death (infarct) are still considered gold standard.^5^ Since this technique requires sectioning and staining of ischemic brain tissue (using 2,3,4-triphenyl tetrazolium chloride or Nissl ^6,7,8^), there is a significant concern that tissue distortion may interfere with visualization of the infarcted tissue. Histological processing can introduce changes in brain morphology (swelling/shrinkage) that affects the accuracy of lesion size measurements.^9^ Additionally, histological staining is manual-labor-intensive and can be subject to variability in staining intensity and interpretation leading to errors in lesion size quantification. The choice of the staining methods or the criteria used to define the lesion border can make it difficult to compare ischemic area across different studies. Moreover, there is a whole range of different calculations/estimates that are used to determine infarct area and volume. It can be done using software (e.g. FIJI software^10^) which might allow automatic infarct volume determination.^11^ Otherwise, one can also manually identify regions of interest and lesion volumes can then be calculated as the sum of the sectional infarct areas multiplied by the interval thickness.^11^

With the development of specialized small animal scanners, magnetic resonance imaging (MRI) has more recently become a promising modality in preclinical laboratories that provides high-resolution images allowing for accurate and precise measurements of stroke size.^12^ Additionally, since MRI is commonly employed in clinical practice, it is a valuable tool for translating preclinical research findings. But since MRI equipment and maintenance can be expensive, its accessibility in preclinical research is still limited. There are several protocols in development to estimate lesion volumes based on *ex vivo* MRI data, however, most require advanced expertise and commercial softwares.^13^

The goal of the present study was to develop a straightforward semi-automated toolkit to estimate lesion volumes of ischemic mouse brains that can be applied to both whole brain scans and brain sections. MRI was performed on formalin-fixed whole mouse brains from a photothrombotic model of stroke followed by macroscopic histological evaluation. We further set out to compare, within the same animals, infarct areas and volumes quantified from post-mortem MRI with that obtained from histological sections stained with Nissl. This study shows that using our toolkit, MRI and histological evaluation both yield to comparable stroke volumes and can be used interchangeably depending on the individual experimental setup.

## Results

We developed a comprehensive step-by-step guide that offers a reliable method for accurately determining stroke volume based on data acquired by post-mortem MRI and histology (**Fig. 1**).

**Fig. 1:**
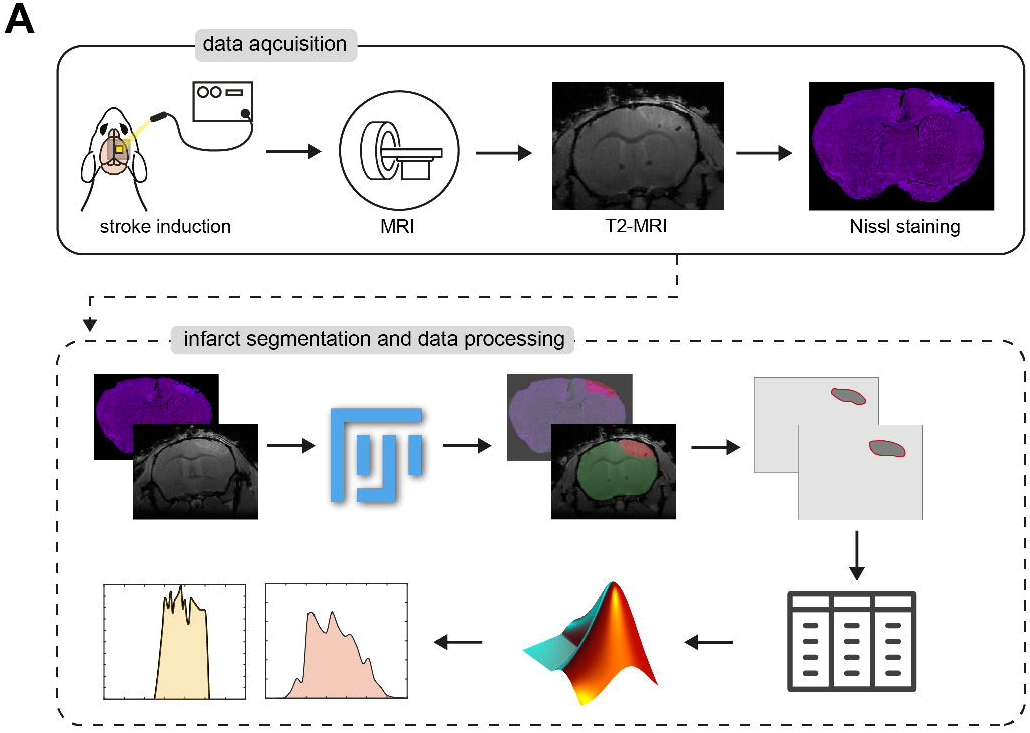
Pipeline for infarct volume estimation based on *ex vivo* MRI and histology. Data acquisition (upper panel): The photothrombotic model was used for infarct induction followed by post-mortem MRI of whole heads. After MRI, brains were removed, sectioned, and histologically processed using fluorescence Nissl stain. Data processing (lower panel): Lesioned areas were identified on all MRI scans and Nissl-images using Fiji (ImageJ). Infarct regions of interest (ROIs) were extracted and referenced to the mouse brain atlas. Areas of all cross sections were measured using MATLAB software followed by volume quantification.

We induced a photothrombotic stroke in the right sensorimotor cortex in C57BL/6 mice (**Fig. 2A**). Successful stroke induction was verified using Laser Doppler imaging (LDI, **Fig. 2B**) confirming severe cortical blood flow reduction of more than 70% compared to baseline levels in the lesioned hemisphere directly after surgery (acute = −71.8%, chronic = −71.7%, p=0.98, **Fig. 2C**, Suppl. Table 1). Brain tissue was collected at two different time points: acutely at 3 days post injury (dpi) and at chronic stages, 28 dpi. After perfusion, whole heads were preserved in 4%-paraformaldehyde (PFA) for 36 hours in preparation for the *ex vivo* MRI. After MRI acquisition, brains were removed and sliced into coronal sections to undergo histological processing.

**Fig. 2.**
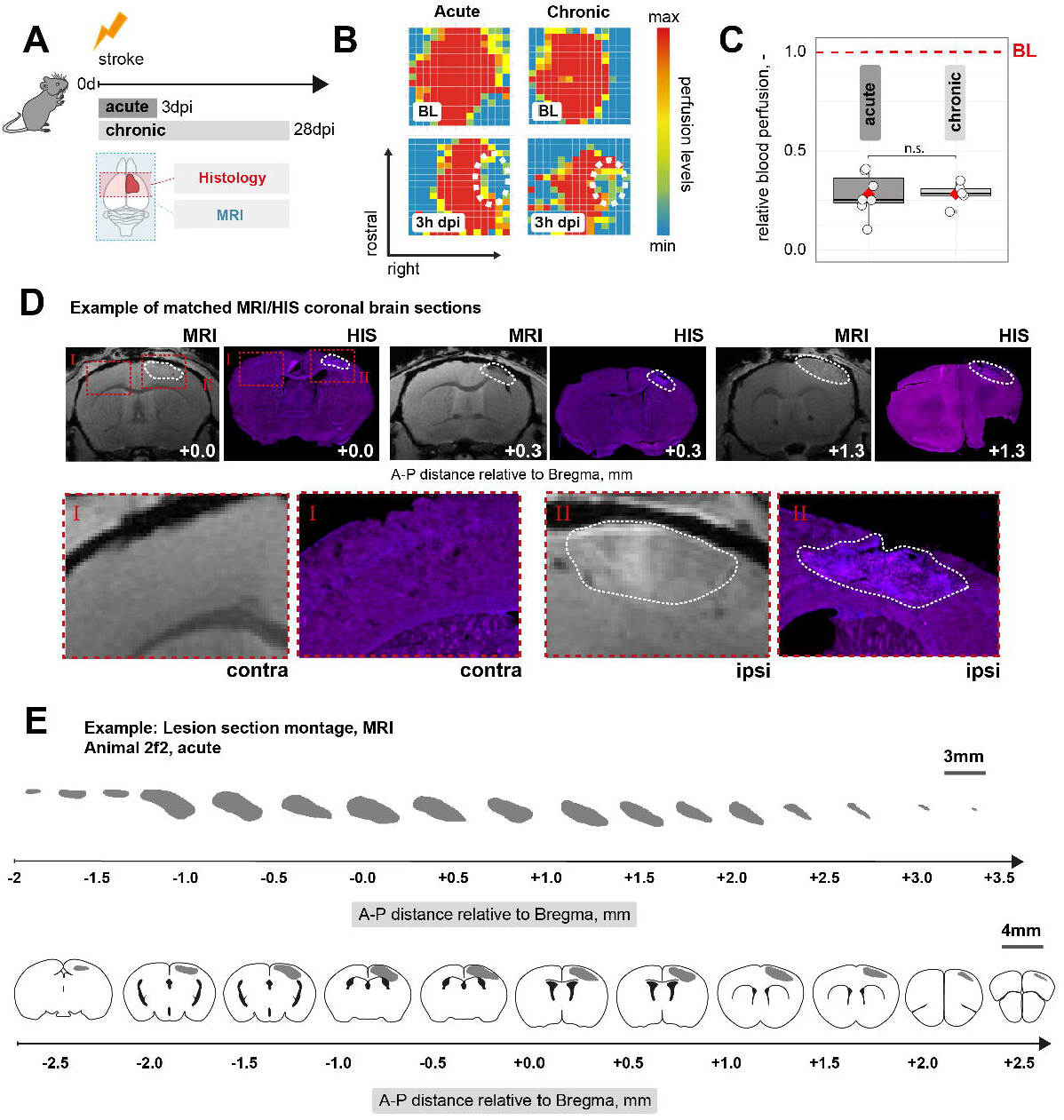
Stroke induction, validation and referencing to brain atlas. (A) Schematic representation of experimental set-up and groups of mice. (B) Representative images of relative blood perfusion at baseline and 3 h after stroke induction. (C) Quantification of cerebral blood perfusion of the injured hemisphere at 3 h following injury relative to baseline blood perfusion. (D) Representative images show ischemic lesion (white outline) on MRI and Nissl stained coronal brain sections at 3 days after injury. (E) Example of manually outlined and extracted infarct ROIs. The ROIs were referenced to the mouse brain atlas according to their anterior-posterior (A-P) distance to Bregma. Scale bar: 3 mm (top) and 4 mm (bottom). Boxplots indicate the 25% to 75% quartiles of the data. For boxplots: each dot in the plots represents one animal. Significance of mean differences between the groups (acute and chronic) was assessed using unpaired two-sample t-test. BL, baseline; dpi, days post injury; HIS, histology; MRI, magnetic resonance imaging; contra, contralesional; ipsi, ipsilesional, ROI: region of interest

Stroked areas were calculated based on MRI-acquired images and histologically processed (Nissl-stained) sections. Sections through the major areas of the infarct (in the range -2mm to +3.5mm relative to bregma, in anterior-posterior direction) were matched (i.e., MRI images with the processed Nissl images, **Fig. 2D**). The infarcted regions were identified by abnormal signal intensity and the clear contrast between the normal and the infarcted, atrophic zone. The hyperintense morphological changes seen in the T2-MRI are indicative of brain swelling and increased water content in the stroked tissue. Brain section stained with fluorescent Nissl-reveal shrunken and distorted neurons. Lesioned areas were manually outlined on each section using Fiji (ImageJ). Each outlined infarct section was extracted and referenced to the mouse brain atlas according to their anterior-posterior distance to bregma (**Fig. 2E**).^14^

Areas of all cross-sections (MRI and Nissl-stained images) were measured and analyzed using MATLAB. To interpolate data within the range of our measurements we used the makima (**M**odified **A**kima **M**ethod with **I**mproved **A**ccuracy) function.^14^ No significant differences were observed between MRI analysis and histological analysis in quantifying lesion area for both the acute and chronic timepoint. However, MRI may provide a higher sensitivity in identifying subtle changes in tissue integrity, especially those associated with small infarct areas in the most rostral and caudal areas of the stroke, which may not be detected with histology (**Fig. 3A, B**).

**Fig 3.**
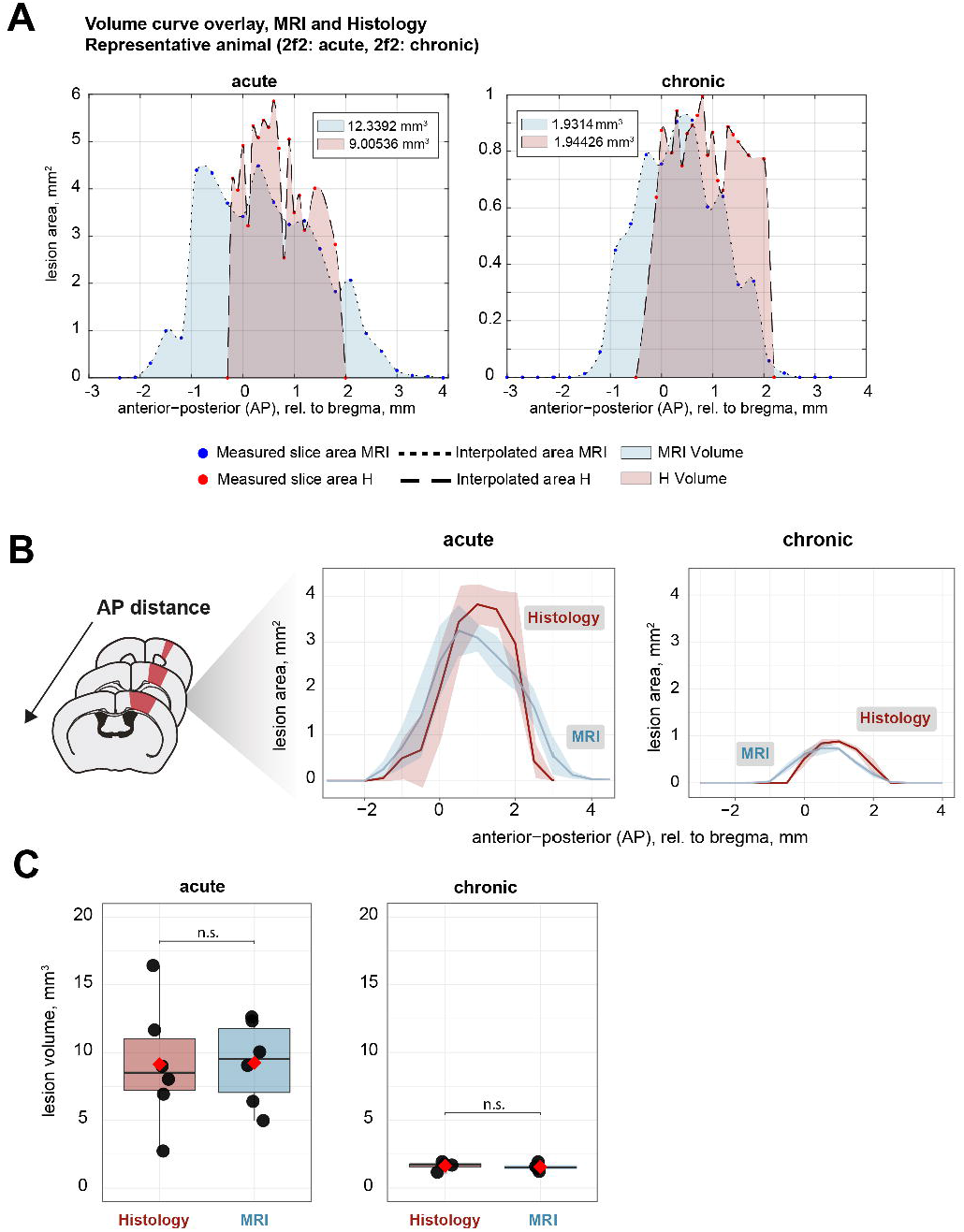
Quantification of infarct volumes from MRI of whole brain tissue and histological analysis of sliced brain sections. (A) Example of two volume curve overlays of both approaches (MRI and histology) at 3 and 28 days after stroke induction. (B) Quantification of lesion area at day 3 and 28 after stroke induction based on MRI and histology. (C) Quantification of lesion volume at day 3 and 28 after stroke induction based on MRI and histology. Data are shown as mean distributions where the red dot represents the mean. Boxplots indicate the 25% to 75% quartiles of the data. For boxplots: each dot in the plots represents one animal. Significance of mean differences between the groups (histology and MRI) was assessed using unpaired two-sample t-test. MRI, magnetic resonance imaging; H, histology.

Next, total stroke volumes were calculated from the area under the curve (AUC) using a trapezoidal computation (Fig 3A, B). No significant difference could be found between MRI-based analysis and histology for both, acute (MRI: 9.24 ± 3.09 mm^3^; histology: 9.14 ± 4.61 mm^3^, p = 0.97) and chronic (MRI: 1.56 ± 0.29 mm^3^; histology: 1.64 ± 0.33 mm^3^, p=0.73, Suppl. Table 1) timepoints.

Previous reports have shown that stroke size decreases between the initial (acute) and long-term (chronic) stages largely as a result of tissue distortion, phagocytosis, resolution of edema, glial and neural remodeling ^15^. In line with expectations, we observe a comparable decrease in infarct volume between the acute (3dpi) and chronic (28dpi) phase (MRI = -83.2% and histology: - 81.2%).^16,17^

## Discussion

In preclinical research, accurate quantification of the lesion size is crucial; it helps to assess the effectiveness of treatments and to compare results across different studies. Several methods are being used to assess stroke size, with histopathology being considered the gold standard. However, there are certain limitations that come along with this approach: Histology only provides two dimensional images, which can lead to underestimation of the true lesion extent. Secondly, brain tissue processing can cause artifacts, such as shrinkage or distortion, which can further affect the accuracy of the measurements. Additionally, histological methods can be time-consuming and labor-intensive (e.g., sectioning, staining, mounting), making it challenging to process a large number of samples. In the last decades, a vast number of new imaging tools have emerged that allow for a reliable assessment of stroke volumes. One such method is magnetic resonance imaging (MRI). MRI provides high-resolution images allowing for accurate and precise measurements of stroke size. And with the introduction of small animal scanners, it has now become a promising tool commonly used in preclinical research.

In this study, we developed a comprehensive step-by-step guide to quantify stroke volume. We employed two distinct techniques for estimating lesion area and volume in an experimental photothrombotic stroke model: (1) based on data acquired by post-mortem MRI and (2) based on a histological approach, where brain sections stained with fluorescent Nissl reveal shrunken and distorted neurons indicating tissue necrosis. The use of perfused-fixed brains throughout the whole study allowed us to directly compare the results. Both techniques, MRI and histology, were capable of visualizing all parameters of infarction/well-defined ischemic area. And despite the mentioned confounders, such as tissue shrinkage or loss of material from the lesion core during histological processing, we found that the final infarct volume on MRI was very well correlated with the infarct volume measured on Nissl-stained sections.

MRI provides a three-dimensional and high-resolution view of the brain, which is essential for accurate assessment of stroke size. Even though the use of MRI to image small animals is getting more popular, it is still restricted to well-funded research centers.^18,19^ A few studies have been conducted to contrast the effectiveness of MRI and histology in experimental stroke models. In a study analyzing multiparametric MRI data after ischemia in rats, a tissue signature analysis demonstrated a high degree of correlation with the histological score at different timepoints.^20^ In contrast, others reported that negative MRI (DWI and T2WI) findings 72h after ischemia may not indicate normal tissue status as seen in the histological outcome. ^21^ It is possible that the detection of tissue damage by MRI depends on the extent of tissue injury. Partial damage to neurons and glial cells with preservation of tissue structure (known as incomplete infarction) can go unnoticed amidst the seemingly normal cells; and may therefore not be detectable by MRI.^21,22^ It also seems reasonable to assume that the significant variability in MRI protocols, handling and positioning of the animals and postprocessing methods might affect lesion size measurements and therefore complicate the comparison to well-standardized histological measures.^23^

The advantage of *ex-vivo* imaging is that it allows for high-throughput acquisition of multiple sections, providing morphologically detailed data of an entire target organ, while avoiding contamination from physiological sources such as pulse, respiration, swallowing and other movements. The use of *ex vivo* brains can notably increase the image resolution and even facilitate additional morphological analysis. Furthermore, the specimen stays intact for follow-up histology. For instance, sliced brain sections can be stained to analyze histological changes associated with regeneration including inflammation, scarring, neurogenesis and vascular repair ^24,25^.

Histological methods for lesion evaluation usually include the slicing of brain sections followed by either 2,3,5-triphenyltetrazolium chloride (TTC) staining, haematoxylin and eosin (H&E) staining, or cresyl violet staining. However, many studies demonstrate that TTC staining can be accurately performed as late as 7 days after stroke, making it unusable for long-term studies. Another possibility is to immunolabel for specific cellular markers (such as NeuN or MAP2, using fixed tissue). Immunolabeling might give rise to images with greater resolution, even down to cellular and subcellular levels. We used NeuroTrace, a fluorescent Nissl stain, to histologically identify the lesioned area. However, there are several other cell markers that can be applied to locate ischemic regions. Neuronal markers such as NeuN (Neuronal Nuclei), MAP2 (Microtubule-Associated Protein 2) or βIII-Tubulin help to distinguish between healthy and damaged neurons. Reactive astrocytes and activated microglia can be identified using GFAP (Glial Fibrillary Acidic Protein) or Iba1 (Ionized calcium-binding adapter molecule 1). These markers are used to locate the glial scar and the ischemic core, respectively. CD31 is a vascular marker that reflects ischemia-induce changes in vasculature, such as endothelial cell swelling or blood brain barrier (BBB) leakage.^26,24^ Summing up, the selection of specific markers depends on the research question and the nature of the stroke model being studied.

Preclinical studies emphasize the validity of lesion volume quantification to evaluate the effectiveness of potential treatments for stroke. E.g., cell-based therapy has been demonstrated to reduce infarct volume while improving neurological function deficits in different experimental stroke models. The systemic application of mesenchymal stem cells (MSCs) significantly decreases lesion volumes in different experimental stroke models and the intravenous transplantation of neural progenitor cells (NPCs) leads to a reduction of infarct size along with long-term functional amelioration after ischemia.^27,28,29^ Antibody-based treatment strategies have also been proven to effectively reduce infarct volumes and improve neuronal performance in different experimental stroke models.^30,31^ Inhibition of TRL4 using monoclonal antibodies decreased both infarct volume and brain swelling in MCAO mice compared to an untreated group.^32^ Likewise, approaches to regulate angiogenesis (so called pro- and antiangiogenic therapies) are increasingly being explored.^33^ Ventricular injections of VEGF have been shown to stimulate angiogenesis and reduce infarct volume in adult rats and the genetic overexpression of VEGF increased brain microvessel density after MCAO in mice.^34,35^

Infarct size is often used as a surrogate measure for functional outcome and has been shown to correlate with e.g. the Bederson score, the forelimb placing performance, the water maze test or overall sensorimotor asymmetry quantification.^36,37,38^ However, other attempts to correlate behavior outcomes with infarct volumes have met mixed results, questioning the validity of the assumption that a larger stroke corresponds to more severe neurological deficits.^39,40^ Consistency in lesion measurement techniques and protocols would allow for more reliable conclusions. Standardization could enable the pooling of data from multiple studies and shed light on the likelihood of volume-function correlations in preclinical research.

The combination of *ex vivo* MRI and histopathology should be considered, not only for analyzing lesion dimensions but also for other neurological research questions. Due to the brain’s heterogeneity, histological sectioning often fails to provide a complete view of every region. MRI on a fixed brain could guide pathologists to sample specific areas for subsequent work-up.^41^ For stem cell applications, preclinical assessments are required to test for potential tumor induction. *Ex vivo* MRI can be used to scan the brain and the spinal cord following cell injection to identify specific (abnormal pale) areas, which potentially correlate histopathologically to teratomas.^42^ Using both MRI paired with gross histological will optimize time of sacrifice and selection of an appropriate stain and improve the scientific significance of the experimental data.

The photothrombotic model of stroke is a well-standardized model that has many strengths, including consistent location and size of infarct, relatively simple surgical procedures, and a low mortality rate.^43,44^ Despite being a rather mild model compared to other experimental stroke models such as MCAO, we found a robust loss of neurons in the infarcted regions and well-defined ischemic border zones. This is consistent with own findings from previous studies, in which we additionally showed that neuronal loss is usually accompanied by fewer astrocytes and decreased microglia activation in the stroke core zone.^45,26^ The lesion volumes observed in this study are comparable to previous studies that have used the photothrombotic stroke model.^46^ There are various other experimental methods used to induce focal ischemia which result in more extensive cortical damage. Middle cerebral artery occlusion (MCAO) typically affects a large portion of the brain and can cause severe neurological deficits. However, size and location of the infarct can be variable, making it more difficult to standardize compared to the photothrombotic model.^47^

In conclusion, we describe a toolkit that allows straightforward lesion volume estimation of ischemic mouse brains. This toolkit generates results that are highly correlated when assessing stroke volumes, whether using full brain MRI images or sliced Nissl-stained brain sections at acute and chronic time points after stroke.

## Methods & Materials

### Study design

Methods for measuring lesion volume were compared utilizing histology and MRI from the brains of mice (n=10) that had undergone cortical ischemia. All animals received a large photothrombotic stroke to the right sensorimotor cortex. At 3 and 28 days after injury induction, whole heads were collected, formalin-fixed and imaged using T2-weighted diffusion MRI. Brains were removed, dissected, and histologically analyzed. We chose the timepoints based on previous literature^48^; animals were categorized according to the phase of stroke as acute (<7 days post-stroke) or chronic (≥21 days post-stroke).

### Animals

All procedures were conducted in accordance with governmental, institutional (University of Zurich), and ARRIVE guidelines and had been approved by the Veterinarian Office of the Canton of Zurich. In total, 10 adult female C57BL/6 mice (acute timepoint: 6, chronic timepoint: 4) were used. Breeding of C57BL/6 mice was performed at Laboratory Animal Services Center (LASC) in Schlieren, CH. All animals were housed in standard type II/III cages on a 12h day/light cycle (6:00 A.M. lights on) with food and water ad libitum. All mice were acclimatized for at least a week to environmental conditions before set into experiment.

### Photothrombotic stroke induction

Anesthesia was performed using isoflurane (5% induction, 1.5-2% maintenance, Attane, Provet AG) and adequate sedation was confirmed by tail pinch. Novalgin (1mg/ml) was applied via drinking water; 24 h prior to the procedure and for three consecutive days directly after stroke surgery. Cerebral ischemia was induced by photothrombotic stroke surgery as previously described.^24,31,45,49–51^ Briefly, animals were fixed in a stereotactic frame (David Kopf Instruments), the surgical area was sanitized using betadine (Mundipharma, Germany), and the skull was exposed through a cut along the midline. A cold light source (Olympus KL 1,500LCS, 150W, 3,020K) was positioned over the right forebrain cortex (anterior/posterior: −1.5–+1.5 mm and medial/lateral 0 mm to +2 mm relative to Bregma). Rose Bengal (15 mg/ml, in 0.9% NaCl, Sigma) was injected intraperitoneally 5 min prior to illumination and the region of interest was subsequently illuminated through the intact skull for 10 min. To restrict the illuminated area, an opaque template with an opening of 3 × 4 mm was placed directly on the skull. The wound was closed using a 6/0 silk suture and animals were allowed to recover.

### Blood perfusion by Laser Doppler imaging

Cortical perfusion was evaluated using Laser Doppler Imaging (Moor Instruments, MOORLDI2-IR). Briefly, animals were fixed in a stereotactic frame and the region of interest was shaved and sanitized. A cut was made along the midline to uncover the skull, and the brain was scanned using the *repeat image measurement mode*. The resulting data was exported and quantified using Fiji (Image J) in terms of total flux in the ROI.

### Sample preparation and diffusion protocol for MRI

Animals were euthanized using pentobarbital (i.p, 150 mg/kg body weight, Streuli Pharma AG) and transcardially perfused with Ringer solution (containing 5 ml/l Heparin, B. Braun) followed by paraformaldehyde (PFA, 4%, in 0.2 M phosphate buffer, pH 7). For MRI procedure, whole mouse heads were collected and post-fixed in 4% paraformaldehyde (PFA) solution for 36 hours. MRI was performed on a 7T small animal scanner with 16 cm bore size (Bruker, Ettlingen, Germany). The fixed brains were put into an Eppendorf cap and imaged using a cryogenically cooled quadrature surface coil (Cryocoil, Bruker, Fällanden, Switzerland). A package of 20 slices with 0.3 mm thickness (no interslice gap) was acquired with a FLASH sequence with a field of view of 15 mm x 15 mm and matrix size of 300 × 240, yielding a spatial in-plane resolution of 50 μm x 50 μm (echo time TE=10 ms, repetition time TR=400 ms, 10 repetitions, total scan time 10 min 40 s).

### Histology

After MRI acquisition, brains were removed and transferred to 30% sucrose for 3 consecutive days for cryoprotection. Coronal sections were cut (40um, Microm HM430, Leica) and kept as free-floating sections in a cryoprotectant solution (PBS, ethylene glycol, sucrose) at -20° C. For immunostainings, sections were blocked with 5% normal donkey serum for 1h at room temperature and incubated with Nissl (1:2000, in 0.1M PB, Sigma) for 30 minutes. Sections were mounted using Mowiol.

### Data processing

The estimation of lesion volumes was divided into 3 stages. In the first stage, the lesion area was manually traced/outlined on each coronal brain section (MRI images and Nissl-stained images) using FIJI (*ImageJ*, version 2.1.0/1.53c). In the second stage, once regions of the lesion have been identified, numerical values for the number of pixels in the selection were obtained and converted into mm^2^. In the third stage, these values were imported into MATLAB (R2022a, The Mathworks, Natick, MA, USA) and lesion volumes were calculated using a customized script. For step-by-step guidance, please refer to our protocol (Supplementary Materials).

### Image pre-processing

Nissl-stained sections were visualized using an Axio Scan Z.1 slide scanner (Carl Zeiss, Germany) with a 20x/0.8 objective lens and later processed and exported as -.tiff files using the ZenBlue software (Version 3.5, Carl Zeiss, Germany).

#### Lesion area measurement

The infarcted areas were estimated by a single blinded researcher using the software FIJI (ImageJ, version 2.1.0/1.53c). First, the images obtained from the MRI were anatomically matched with the sections that were histologically processed. On each brain section (MRI and Nissl-stained), the ischemic area was visually identified as defined by the area with atypical and atrophic tissue morphology including pale areas with lost Nissl staining. The identified ischemic area was then outlined using the freehand polygon tool. All outlined ROIs were measured (numerical value for the number of pixels) and converted to mm^2^ using an index ruler of known size which was included beforehand. Values were stored in a .-csv table.

#### Volume estimation

Once the lesion area in each slice was estimated, the lesion volume was calculated using a customized MATLAB script (see Suppl. Materials). We used the *modified akima interpolation* (makima) function to interpolate the missing values. The area under the curve (AUC) was calculated using the trapezoidal computation rule. Briefly, the volume was calculated by multiplication of the lesion area and the distance between the sections as follows:

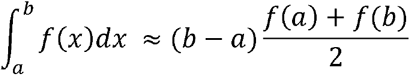

*f*(*x*) represents the cross-sectional area at *x*.

### Statistics

Statistical analysis was performed using R-Studio. Sample sizes were designed with adequate power according to previous studies. All data were tested for normal distribution by using the Shapiro-Wilk test. The significance of mean differences between normally distributed data (MRI vs. Histology) were tested for differences with a two-tailed unpaired one-sample t-test. Data are expressed as means ± SD, and statistical significance was defined as *p < 0.05, **p < 0.01, and ***p < 0.001.

## Supporting information

Suppl. Protocol

## Data availability

The software code and raw data utilized in this study can be accessed in the Supplementary Material and are also available online through Zenodo: 10.5281/zenodo.8056094

## Acknowledgements

-

## Author contributions

RZW, DB, MA, DR, CT, RR designed the study, prepared the figures, and wrote the manuscript. RZW, NHR, SH, BAB and MA carried out the experiments. All authors read and approved the final manuscript.

## Declaration of interests

The authors declare that the research was conducted in the absence of any commercial or financial relationships that could be construed as a potential conflict of interest.

## Notes

https://zenodo.org/record/8056094

### Competing Interest Statement

The authors have declared no competing interest.

## References

1. Heart Disease and Stroke Statistics—2016 Update | Circulation. Accessed April 18, 2023. https://www.ahajournals.org/doi/10.1161/CIR.0000000000000350?url_ver=Z39.88-2003&rfr_id=ori:rid:crossref.org&rfr_dat=cr_pub%20%200pubmed

2. McDermott M, Skolarus LE, Burke JF. A systematic review and meta-analysis of interventions to increase stroke thrombolysis. BMC Neurol. 2019;19(19):86. doi:10.1186/s12883-019-1298-2

3. Lourbopoulos A, Mourouzis I, Xinaris C, et al. Translational Block in Stroke: A Constructive and “Out-of-the-Box” Reappraisal. Front Neurosci. 2021;15:652403. doi:10.3389/fnins.2021.652403

4. Ludewig P, Gdaniec N, Sedlacik J, et al. Magnetic Particle Imaging for Real-Time Perfusion Imaging in Acute Stroke. ACS Nano. 2017;11(11):10480–10488. doi:10.1021/acsnano.7b05784

5. Mulder IA, Khmelinskii A, Dzyubachyk O, et al. Automated Ischemic Lesion Segmentation in MRI Mouse Brain Data after Transient Middle Cerebral Artery Occlusion. Front Neuroinformatics. 2017;11. Accessed April 18, 2023. https://www.frontiersin.org/articles/10.3389/fninf.2017.00003

6. García-Cabezas MÁ, John YJ, Barbas H, Zikopoulos B. Distinction of Neurons, Glia and Endothelial Cells in the Cerebral Cortex: An Algorithm Based on Cytological Features. Front Neuroanat. 2016;10. Accessed April 18, 2023. https://www.frontiersin.org/articles/10.3389/fnana.2016.00107

7. Atochin DN, Chernysheva GA, Aliev OI, et al. An improved three-vessel occlusion model of global cerebral ischemia in rats. Brain Res Bull. 2017;132:213–221. doi:10.1016/j.brainresbull.2017.06.005

8. Choi CH, Yi KS, Lee SR, et al. A novel voxel-wise lesion segmentation technique on 3.0-T diffusion MRI of hyperacute focal cerebral ischemia at 1⍰h after permanent MCAO in rats. J Cereb Blood Flow Metab. 2018;38(38):1371–1383. doi:10.1177/0271678X17714179

9. Bay V, Iversen NK, Shiadeh SMJ, Tasker RA, Wegener G, Ardalan M. Tissue processing and optimal visualization of cerebral infarcts following sub-acute focal ischemia in rats. J Chem Neuroanat. 2021;118:102034. doi:10.1016/j.jchemneu.2021.102034

10. Schindelin J, Arganda-Carreras I, Frise E, et al. Fiji: an open-source platform for biological-image analysis. Nat Methods. 2012;9(9):676–682. doi:10.1038/nmeth.2019

11. Sommer C. Histology and Infarct Volume Determination. In: Dirnagl U, ed. Rodent Models of Stroke. Neuromethods. Humana Press; 2010:213–226. doi:10.1007/978-1-60761-750-1_15

12. Raylman RR, Ledden P, Stolin AV, Hou B, Jaliparthi G, Martone PF. Small animal, positron emission tomography-magnetic resonance imaging system based on a clinical magnetic resonance imaging scanner: evaluation of basic imaging performance. J Med Imaging. 2018;5(5):033504. doi:10.1117/1.JMI.5.3.033504

13. Gabrielson K, Maronpot R, Monette S, et al. In Vivo Imaging With Confirmation by Histopathology for Increased Rigor and Reproducibility in Translational Research: A Review of Examples, Options, and Resources. ILAR J. 2018;59(59):80–98. doi:10.1093/ilar/ily010

14. Modified Akima piecewise cubic Hermite interpolation - MATLAB makima. Accessed April 18, 2023. https://www.mathworks.com/help/matlab/ref/makima.html

15. Gaudinski MR, Henning EC, Miracle A, Luby M, Warach S, Latour LL. Establishing Final Infarct Volume. Stroke. 2008;39(39):2765–2768. doi:10.1161/STROKEAHA.107.512269

16. Li H, Zhang N, Lin HY, et al. Histological, cellular and behavioral assessments of stroke outcomes after photothrombosis-induced ischemia in adult mice. BMC Neurosci. 2014;15(15):58. doi:10.1186/1471-2202-15-58

17. Shen F, Jiang L, Han F, Degos V, Chen S, Su H. Increased Inflammatory Response in Old Mice is Associated with More Severe Neuronal Injury at the Acute Stage of Ischemic Stroke. Aging Dis. 2019;10(10):12–22. doi:10.14336/AD.2018.0205

18. Xie L, Cianciolo R, Hulette B, et al. Magnetic Resonance Histology of Age-related Nephropathy in the Sprague Dawley Rat. Toxicol Pathol. 2012;40(40):764–778. doi:10.1177/0192623312441408

19. Johnson GA, Badea A, Jiang Y. Quantitative neuromorphometry using magnetic resonance histology. Toxicol Pathol. 2011;39(39):85–91. doi:10.1177/0192623310389622

20. Jacobs MA, Zhang ZG, Knight RA, et al. A model for multiparametric mri tissue characterization in experimental cerebral ischemia with histological validation in rat: part 1. Stroke. 2001;32(32):943–949. doi:10.1161/01.str.32.4.943

21. Li F, Liu KF, Silva MD, et al. Transient and permanent resolution of ischemic lesions on diffusion-weighted imaging after brief periods of focal ischemia in ratslll: correlation with histopathology. Stroke. 2000;31(31):946–954. doi:10.1161/01.str.31.4.946

22. Garcia JH, Liu KF, Ye ZR, Gutierrez JA. Incomplete infarct and delayed neuronal death after transient middle cerebral artery occlusion in rats. Stroke. 1997;28(28):2303-2309; discussion 2310. doi:10.1161/01.str.28.11.2303

23. Milidonis X, Marshall I, Macleod MR, Sena ES. Magnetic Resonance Imaging in Experimental Stroke and Comparison With Histology. Stroke. 2015;46(46):843–851. doi:10.1161/STROKEAHA.114.007560

24. Rust R, Grönnert L, Gantner C, et al. Nogo-A targeted therapy promotes vascular repair and functional recovery following stroke. Proc Natl Acad Sci. 2019;116(116):14270–14279. doi:10.1073/pnas.1905309116

25. Rust R, Kirabali T, Grönnert L, et al. A Practical Guide to the Automated Analysis of Vascular Growth, Maturation and Injury in the Brain. Front Neurosci. 2020;14. doi:10.3389/fnins.2020.00244

26. Weber RZ, Mulders G, Perron P, Tackenberg C, Rust R. Molecular and anatomical roadmap of stroke pathology in immunodeficient mice. Front Immunol. 2022;13. Accessed April 19, 2023. https://www.frontiersin.org/articles/10.3389/fimmu.2022.1080482

27. Bacigaluppi M, Russo GL, Peruzzotti-Jametti L, et al. Neural Stem Cell Transplantation Induces Stroke Recovery by Upregulating Glutamate Transporter GLT-1 in Astrocytes. J Neurosci. 2016;36(36):10529–10544. doi:10.1523/JNEUROSCI.1643-16.2016

28. Bonsack B, Corey S, Shear A, et al. Mesenchymal stem cell therapy alleviates the neuroinflammation associated with acquired brain injury. CNS Neurosci Ther. 2020;26(26):603–615. doi:10.1111/cns.13378

29. Chung JW, Chang WH, Bang OY, et al. Efficacy and Safety of Intravenous Mesenchymal Stem Cells for Ischemic Stroke. Neurology. 2021;96(96):e1012–e1023. doi:10.1212/WNL.0000000000011440

30. Woods D, Jiang Q, Chu XP. Monoclonal antibody as an emerging therapy for acute ischemic stroke. Int J Physiol Pathophysiol Pharmacol. 2020;12(12):95–106.

31. Rust R, Weber RZ, Grönnert L, et al. Anti-Nogo-A antibodies prevent vascular leakage and act as pro-angiogenic factors following stroke. Sci Rep. 2019;9(9):1–10. doi:10.1038/s41598-019-56634-1

32. Andresen L, Theodorou K, Grünewald S, et al. Evaluation of the Therapeutic Potential of Anti-TLR4-Antibody MTS510 in Experimental Stroke and Significance of Different Routes of Application. PLOS ONE. 2016;11(11):e0148428. doi:10.1371/journal.pone.0148428

33. Rust R, Gantner C, Schwab ME. Pro- and antiangiogenic therapies: current status and clinical implications. FASEB J. 2019;33(33):34–48. doi:10.1096/fj.201800640RR

34. Sun Y, Jin K, Xie L, et al. VEGF-induced neuroprotection, neurogenesis, and angiogenesis after focal cerebral ischemia. J Clin Invest. 2003;111(111):1843–1851. doi:10.1172/JCI17977

35. Wang Y, Kilic E, Kilic U, et al. VEGF overexpression induces post-ischaemic neuroprotection, but facilitates haemodynamic steal phenomena. Brain J Neurol. 2005;128(Pt 1):52–63. doi:10.1093/brain/awh325

36. Bieber M, Gronewold J, Scharf AC, et al. Validity and Reliability of Neurological Scores in Mice Exposed to Middle Cerebral Artery Occlusion. Stroke. 2019;50(50):2875–2882. doi:10.1161/STROKEAHA.119.026652

37. C Turner R, DiPasquale K, F Logsdon A, et al. The role for infarct volume as a surrogate measure of functional outcome following ischemic stroke. J Syst Integr Neurosci. 2016;2(4). doi:10.15761/JSIN.1000136

38. Machado AG, Baker KB, Schuster D, Butler RS, Rezai A. Chronic electrical stimulation of the contralesional lateral cerebellar nucleus enhances recovery of motor function after cerebral ischemia in rats. Brain Res. 2009;1280:107–116. doi:10.1016/j.brainres.2009.05.007

39. Wakayama K, Shimamura M, Sata M, et al. Quantitative measurement of neurological deficit after mild (30 min) transient middle cerebral artery occlusion in rats. Brain Res. 2007;1130(1130):181–187. doi:10.1016/j.brainres.2006.10.088

40. Encarnacion A, Horie N, Keren-Gill H, Bliss TM, Steinberg GK, Shamloo M. Long-term behavioral assessment of function in an experimental model for ischemic stroke. J Neurosci Methods. 2011;196(196):247–257. doi:10.1016/j.jneumeth.2011.01.010

41. Hanig J, Paule MG, Ramu J, et al. The use of MRI to assist the section selections for classical pathology assessment of neurotoxicity. Regul Toxicol Pharmacol RTP. 2014;70(70):641–647. doi:10.1016/j.yrtph.2014.09.010

42. Ramot Y, Schiffenbauer YS, Amouyal N, et al. Compact MRI for the detection of teratoma development following intrathecal human embryonic stem cell injection in NOD-SCID mice. Neurotoxicology. 2017;59:27–32. doi:10.1016/j.neuro.2017.01.003

43. Labat-gest V, Tomasi S. Photothrombotic Ischemia: A Minimally Invasive and Reproducible Photochemical Cortical Lesion Model for Mouse Stroke Studies. J Vis Exp JoVE. 2013;(76):50370. doi:10.3791/50370

44. Sun YY, Kuo YM, Chen HR, Short-Miller JC, Smucker MR, Kuan CY. A murine photothrombotic stroke model with an increased fibrin content and improved responses to tPA-lytic treatment. Blood Adv. 2020;4(4):1222–1231. doi:10.1182/bloodadvances.2019000782

45. Weber RZ, Grönnert L, Mulders G, et al. Characterization of the Blood Brain Barrier Disruption in the Photothrombotic Stroke Model. Front Physiol. 2020;11. Accessed April 18, 2023. https://www.frontiersin.org/articles/10.3389/fphys.2020.586226

46. Porritt MJ, Andersson HC, Hou L, et al. Photothrombosis-Induced Infarction of the Mouse Cerebral Cortex Is Not Affected by the Nrf2-Activator Sulforaphane. PLOS ONE. 2012;7(7):e41090. doi:10.1371/journal.pone.0041090

47. Sommer CJ. Ischemic stroke: experimental models and reality. Acta Neuropathol (Berl). 2017;133(133):245–261. doi:10.1007/s00401-017-1667-0

48. Bogaert-Buchmann A, Poittevin M, Po C, et al. Spatial and Temporal MRI Profile of Ischemic Tissue after the Acute Stages of a Permanent Mouse Model of Stroke. Open Neuroimaging J. 2013;7:4–14. doi:10.2174/1874440001307010004

49. Fan Z, Ardicoglu R, Batavia AA, et al. The vascular gene Apold1 is dispensable for normal development but controls angiogenesis under pathological conditions. Angiogenesis. Published online March 18, 2023. doi:10.1007/s10456-023-09870-z

50. Weber RZ, Mulders G, Kaiser J, Tackenberg C, Rust R. Deep learning-based behavioral profiling of rodent stroke recovery. BMC Biol. 2022;20(20):232. doi:10.1186/s12915-022-01434-9

51. Rust R, Weber RZ, Generali M, et al. Xeno-free induced pluripotent stem cell-derived neural progenitor cells for in vivo applications. J Transl Med. 2022;20(20):421. doi:10.1186/s12967-022-03610-5

