## Supplementary material for "Visualization and estimation of stroke infarct volumes in rodents": Suppl. Protocol

**Practical toolkit to estimate stroke volume in the mouse brain**

The estimation of lesion volumes requires the use of the following programs: Zen Blue (Version 3.5, Carl Zeiss, Germany), FIJI (or ImageJ, version 2.1.0/1.53c) and MATLAB (R2022a, The Mathworks, Natick, MA, USA).

1. **Image pre-processing**
   1.1. Use Zen Blue (Version 3.5, Carl Zeiss, Germany) to process your data.
   1.2. Adjust color/brightness/contrast so that the stroked area is clearly visible.
   1.3. Insert a scale bar (…)
   1.4. Export your files (*Method Parameters/Show all…)*
        1.4.1. Adjust the settings (*File type=Tiff, Compression: LZW*, chose *Channel* and *Scene* of your choice).
2. **Area measurement per coronal brain section**
   Note: For further guidance, see video (Supplementary Materials “Video_area_measurement”).
   2.1. Open the image sequence (-.tiff file).
   2.2. Set the scale (*Analyze/Set Scale…)*
        Note: Use the scale bar of known size that was applied before using the Zen blue software.
   2.3. Open the ROI manager (*Analyze/Tools/ROI Manager…).*
   2.4. Select the *freehand selections* tool and outline the damaged area on each section.
   2.5. Add the selected area to the ROI manager (*ROI Manager/Add[t]…*)
   2.6. Select all ROIs and save them as -.zip file (*ROI Manager/More/Save…*). While all ROIs are still selected, measure (*ROI Manager/Measure…*) and save the results in a -.csv table (*File/Save as…).*
   2.7. Use the FIJI macro (“IsolateSections_FijiMacro.jim”) to isolate the outlined stroke areas.
   2.8. Crop (use the *Rectangle* tool, and then *Analyze/Crop…*) the isolated areas.
   2.9. Insert a scale bar (*Analyze/Tools/Scale bar…*).
   2.10. Save as image sequence (*File/Save as/Image Sequence…*).
3. **Data pre-processing**
   3.1. Before running the script, create a reference table (in excel) where you store the variables required. For the protocol used in this experiment, we used the following variables: Experiment number, name of the output folder, timepoint (acute, chronic), frame number of the scan at Bregma = 0.0, bregma values for the image sequence (lowest to highest). The following image displays the format:

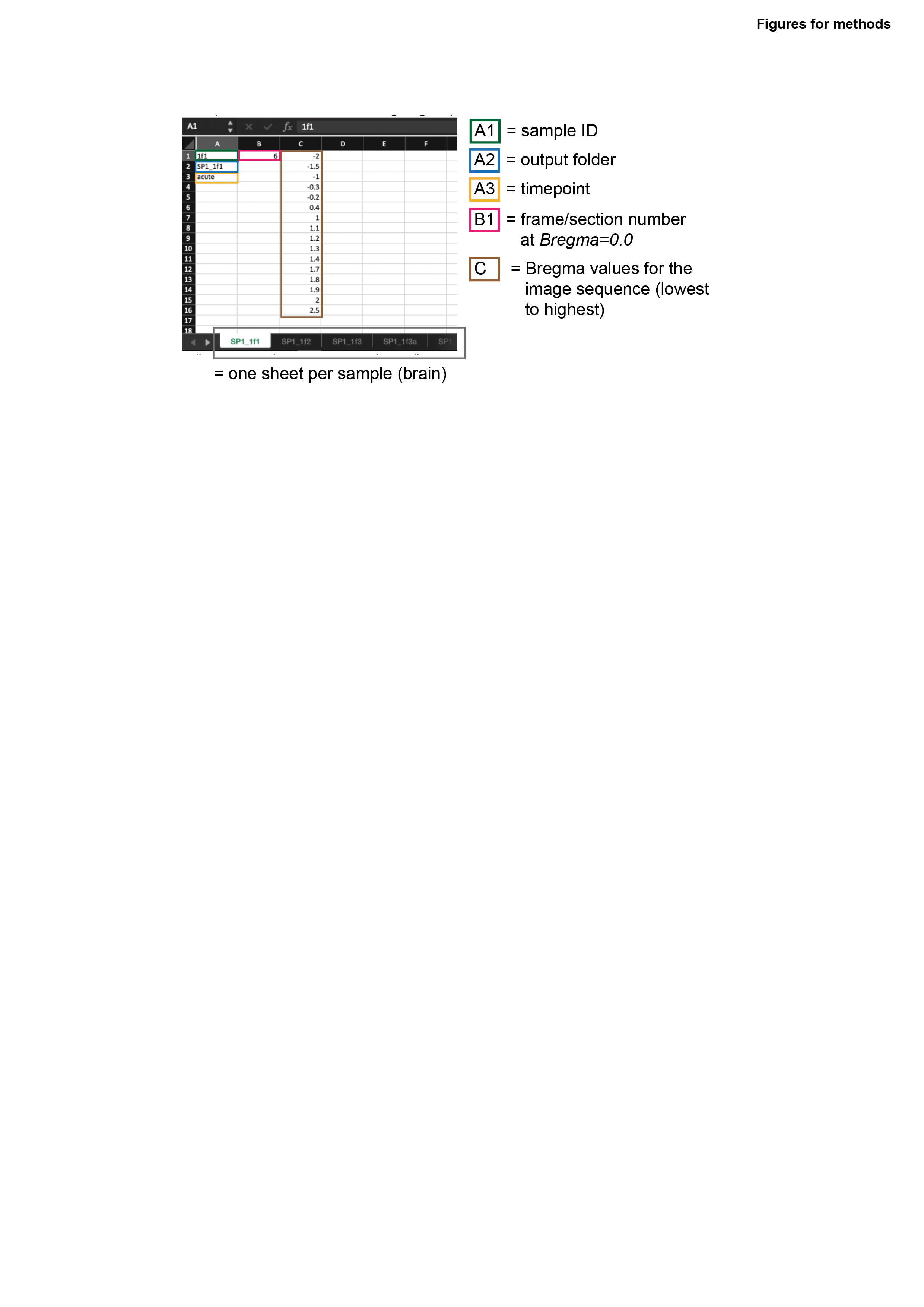

        Each sheet contains a different sample (e.g. brain). Using this format, it is possible to organize different samples into one MATLAB script.
4. **Data processing using MATLAB**
   4.1. Open MATLAB.
   4.2. Import the script (Supplementary material “code_general_single.m”).
   4.3. Select the folder containing all data of all your samples.
        Note: By selecting the data folder containing all your subfolders, the script can run through all folders and access the data files that were exported with FIJI and calculate the volume for every brain simultaneously.
   4.4. When the “*done*” text appears, you may close MATLAB.
        Note: The script has been commented extensively to explain what almost every command line does. This should allow for smooth debugging.
5. **Data post-processing**
   5.1. The MATLAB scripts exports the figures in an editable -.fig format and -.pdf/-.png format. It also exports a -.csv file for every brain.
   5.2. Import the -.csv files into R-Studio for post-processing, plotting and statistical analysis.
